# Extrinsic biological stochasticity and technical noise normalization of single-cell RNA sequencing data

**DOI:** 10.1101/2025.05.11.653373

**Authors:** Meichen Fang, Lior Pachter

## Abstract

The technical noise introduced during single-cell RNA sequencing (scRNA-seq) has led to the use of size factor normalization as a first step prior to data analysis. However, this scaling approach inherently affects extrinsic (between cell) variability of gene expression, which stems from both biological and technical factors. Based on previous models on biological and technical extrinsic noise, we propose a general extrinsic noise model for scRNA-seq to provide a theoretical basis for size factor normalization, thus providing a framework for estimating both biological and technical components of extrinsic noise. We highlight the relationship between normalized gene expression covariance, extrinsic noise, and overdispersion, showing that extrinsic noise explains the baseline overdispersion commonly observed in scRNA-seq data. We validated the technical model by testing the relationship on data from pooled RNA. Interestingly, our model accurately describes mature mRNA counts but not nascent mRNA counts, suggesting the need for an alternative technical model for data derived from nascent transcripts. Using single-cell RNA-seq data, we characterize both biological and technical extrinsic noise and cell size factors estimated using Poisson-like genes. Overall, our model helps clarify common misconceptions and provides insight into the role of extrinsic noise and size factor normalization in scRNA-seq data.

## 1 Introduction

Single-cell RNA sequencing (scRNA-seq) enables genome-wide expression profiling at unprecedented scale, but current data are notoriously noisy, partially due to variability in sequencing depth per cell due to random sampling of libraries during sequencing. This issue becomes particularly acute when the measurement rate is low, and can cause biological signal to be overwhelmed by noise. Therefore, a standard and critical step at the beginning of scRNA-seq analysis is normalization, which is intended to mitigate the effects of technical noise before downstream analysis [1]. Typically, a global-scaling normalization method is used, which calculates a single normalization factor per cell (cell size factor) using the sum of total counts to adjust for variability in sequencing depth and technical artifacts. This approach is implemented in common packages for scRNA-seq analysis [2, 3, 4]. Beyond this, more sophisticated approaches have been developed, including some popular methods that calculate the cell size factor by pooling cells [5], using homogeneously expressed genes [6], introducing multiple cell size factors for different groups of genes [7], and utilizing negative binomial regression [8]. Regardless of the methods used, the common goal of normalization techniques is to use one or more scaling factors to account for technical variation and try to remove it through methods such as scaling and regression.

We argue that common practices for normalization inadvertently remove extrinsic noise. The concept of extrinsic noise emerged first in the study of biological stochasticity in gene expression, and refers to fluctuations in the cellular environment that affect all genes [9]. Applying this concept to scRNA-seq suggests that normalization, particularly scaling, can eliminate biological stochasticity present in extrinsic “noise”. This may be critical as extrinsic noise of biological origin may carry meaningful signals relevant to specific biological questions. As a result, current normalization procedure tend to diminish both biological variation and technical noise [10].

The extrinsic noise in scRNA-seq arises from both biological and technical sources. Previous studies have characterized both types of extrinsic noise and demonstrated that they are prominent. In the context of biological stochasticity, gene expression variability has been classified into extrinsic and intrinsic components based on their underlying mechanisms [9, 11]. Since extrinsic noise affects all genes within a single cell, the normalized covariance between genes has been identified as an effective measure of extrinsic noise in dual-reporter studies [9, 11, 12, 13]. Furthermore, the impact of biological extrinsic noise on transcriptome-wide inference has been explored using a telegraph model, highlighting the importance of accounting for biological extrinsic noise itself, which can also be estimated using normalized covariance between genes and explain the coefficient of variation (CV) of highly expressed genes [14].

Technical noise in scRNA-seq experiments has also been widely studied since the development of single-cell assays. For example, technical noise has been assessed experimentally using ERCC spike-ins to assess technical variance [15, 16, 17]. ERCC-derived technical noise has been linked to global tube-to-tube variations in sequencing efficiency, and a correspondence between technical noise and the observed constant CV for highly expressed transcripts has been noted [16]. UMI counts are typically modeled using binomial or Poisson distributions, corresponding to Bernoulli or Poisson sampling, respectively, with cell-specific capture rates incorporated to account for detection efficiency variability [18, 19, 20, 21]. Notably, in the regime of low capture rates, the Poisson distribution approximates the binomial distribution.

Currently, models for scRNA-seq that account for both biological and technical extrinsic noise are typically based on specific gene expression frameworks and often assume that biological extrinsic noise influences particular kinetic parameters [20, 21]. A more general model that accounts for extrinsic noise from both biological and technical sources without assuming a specific gene expression form could still be valuable. Such a model would allow for flexible characterization and validation of biological intrinsic noise, while generating specific and potentially insightful predictions.

We develop such an extrinsic noise model for scRNA-seq data to account for both biological and technical sources of noise, building on the previous studies described above. We derive a general relationship between observed and intrinsic moments (covariance/variance) under a Bernoulli technical noise model and a scaling assumption for *in vivo* gene expression. In the specific case where genes are independent and exhibit Poisson intrinsic noise, we show that the extrinsic noise is equal to both the normalized covariance and overdispersion. This relationship summarizes two previous results: the estimation of extrinsic noise using normalized covariance originally applied to biological noise [9, 11, 12, 14, 13], and the interpretation of technical variability as a baseline overdispersion observed in pseudocell data [16, 22], generalizing both to total extrinsic noise in scRNA-seq datasets. We test this equality on pooled RNA datasets where counts are intrinsically Poisson-distributed, thereby validating the technical model as well as identifying any abnormalities. Second, when applied to single-cell datasets, this equality enables us to quantify biological and technical extrinsic noise, predict the overdispersion of intrinsically Poisson genes, and identify Poisson genes, whose total expression provides a principled approach for estimating cell size factors. Overall, we demonstrate how a mechanistic model of extrinsic noise clarifies the normalization step in scRNA-seq analysis and offers new insights into the observed variability in scRNA-seq data.

## 2 Results

### 2.1 A single-cell RNA-seq extrinsic noise model

Modeling extrinsic noise in scRNA-seq requires both a biological model of gene expression and a technical model of scRNA-seq measurement so as to jointly account for biological stochasticity and technical noise (Fig 1b). As genes can have very different expression mechanisms and resultant distributions, we do not assume any specific distribution for *in vivo* counts at first. Instead, we only assume that the means of genes in each cell are proportional to a cellular random variable *c*^*bio*^, which represents the cell-wise size factor that summarize the biological extrinsic noise. The value of *c*^*bio*^ could be influenced by many factors such as the cell volume and the cell cycle phase. As there could be many unknown sources of cell-to-cell variability, *c*^*bio*^ is a phenomenological parameter that captures the combined effects of various extrinsic factors. We denote the *in vivo* amount of gene *j* in cell *i* by 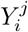 and assume 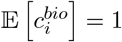 without loss of generality, this model for biological extrinsic noise means

**Figure 1.**
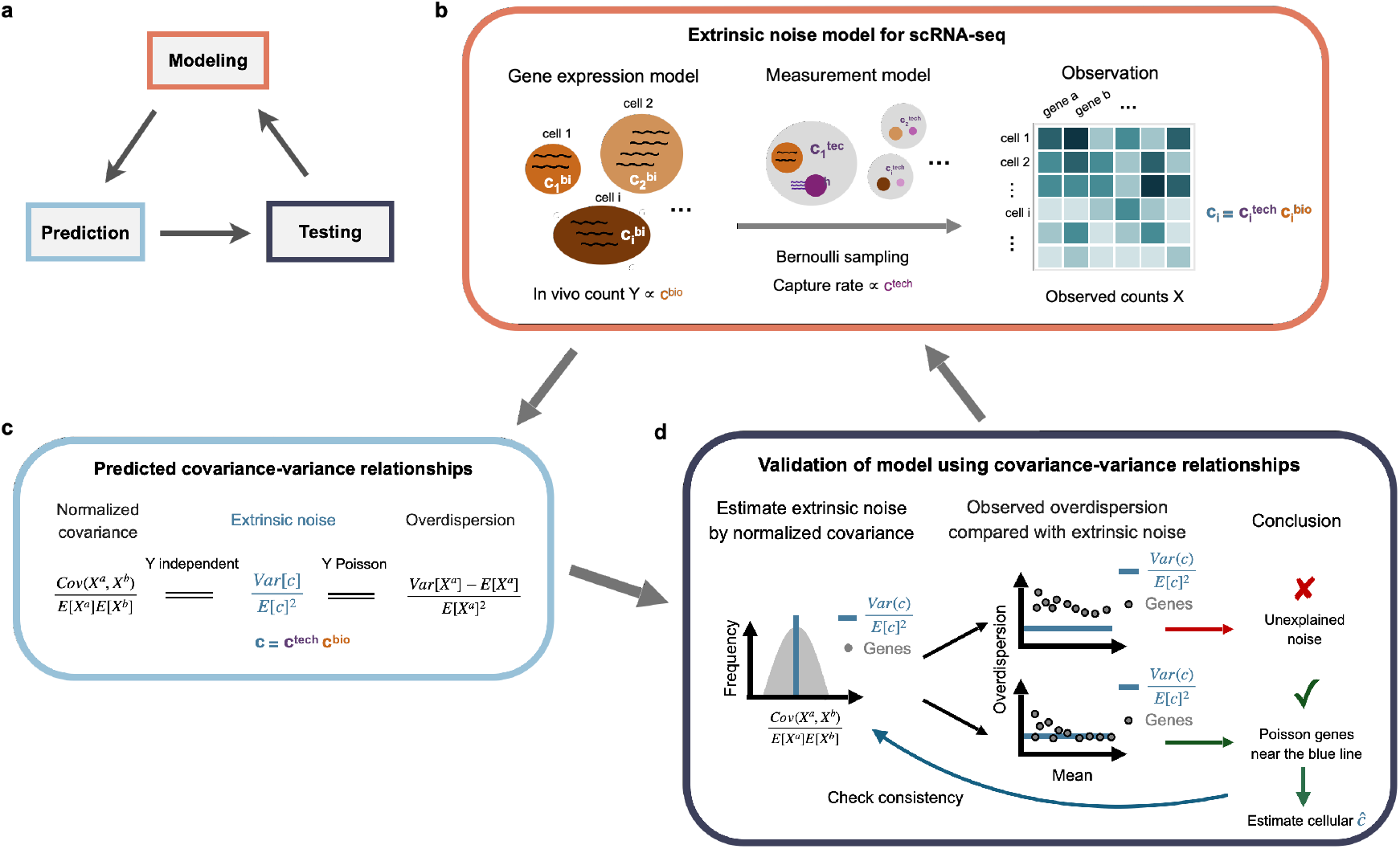
Modeling extrinsic noise in scRNA-seq data. **a)** Model-based closed-loop paradigm. The process begins with the formulation of mechanistic models, followed by rigorous mathematical analysis to generate testable predictions. These predictions are then tested on data, allowing models to be refined or rejected. The cycle repeats with updated models, creating an iterative loop of modeling. **b)** Schematic of the extrinsic noise model. **c)** Predicted relationships among normalized covariance, extrinsic noise, and overdispersion. **d)** Procedure for validating these relationships.

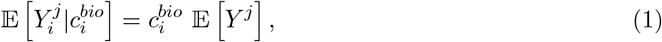

where 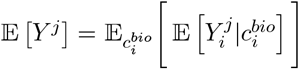 is the mean of gene *j* across cells.

For the technical measurement model, we make much stronger assumptions: we assume a Bernoulli sampling of each transcript in single-cell experiment; this is based on previous studies [22] and the assumptions leads to a binomial distribution of observed counts given *in vivo* counts. This is also the low detection approximation of Poisson sampling (Section 4.1). Similarly, we introduce a cellular random variable *c*^*tech*^ to summarize the relative success probability in the binomial distribution.

This *c*^*tech*^ can be interpreted as affecting relative read depth during sequencing and is independent of the biological model and the *in vivo* counts. To account for differences in capture efficiency between genes, we introduce a constant capture rate, *λ*, as an unknown constant for each molecular species, which cancels out in normalized quantities. Finally, we denote the observed counts after single cell sequencing by *X*, this measurement model yields

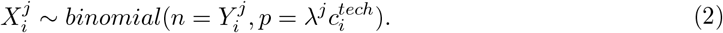

The two assumptions (equation 1 and equation 2) lead to simple expressions that relate the intrinsic normalized (co)variance to the observed normalized (co)variance (Section 4.1):

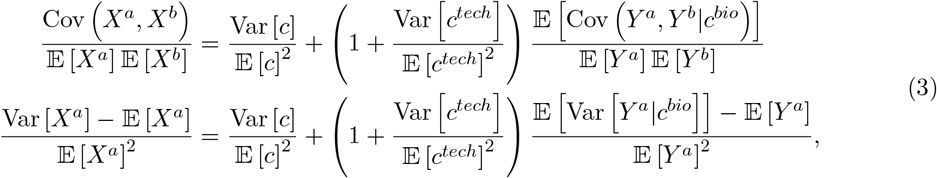

where *a, b* are gene indices and *c* = *c*^*tech*^*c*^*bio*^ is the overall size factor for each cell. We denote 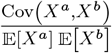 as normalized covariance following previous literature [12], and 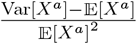 as normalized variance for convenience, since it directly indicates the extent of over-dispersion. The first term on the right hand side 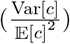 represents the extrinsic noise, representing a combination of both biological and technical extrinsic noise:

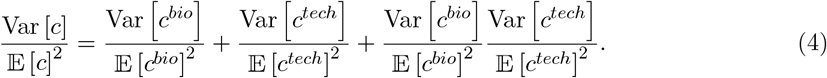

The second terms on the right hand side of equation 3 represent the contribution of intrinsic noise. For example, for two independent and intrinsically Poisson-distributed genes, the intrinsic normalized covariance 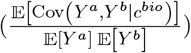 and normalized variance 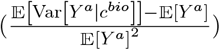 would both be 0. Therefore, the second terms denote the effect of intrinsic normalized (co)variance convoluted with technical extrinsic noise.

The equation 3 leads to two important observations: first, if we assume that genes are intrinsically uncorrelated, the normalized covariance can be used to estimate the extrinsic noise (Fig 1c), which is the canonical approach in previous studies on biological extrinsic noise [11, 12, 14]. Although the exact distribution of normalized covariance between uncorrelated gene pairs depends on the distribution of *c*, it should nevertheless center around the value of extrinsic noise, and we can use the mean/mode of the distribution to estimate the extrinsic noise. If not all but most genes are uncorrelated, the normalized covariance can still provide a reasonable estimate of the extrinsic noise using the mode of the distribution of normalized covariance. The distance between the mode and the mean provides some insight into the correlation between genes since the mode should coincide with the mean if all genes are uncorrelated. Furthermore, if we have an empirical distribution of *c*, we can verify whether the distribution of the normalized covariance aligns with the model (Fig 1d).

Second, if the genes are intrinsically Poisson distributed, then the normalized variance also equals the extrinsic noise (Fig. 1 c). Therefore, after estimating extrinsic noise using normalized covariance between genes, we can test whether each gene is Poisson distributed using this expected equality (Fig. 1 d). Note that the extrinsic noise term in normalized variance contributes to the observed over-dispersion in scRNA-seq data, as it introduces a constant offset visible when plotting the coefficient of variation (CV) against the mean. Therefore, genes are intrinsically less over-dispersed than observed counts might suggest, and it is possible for some genes to not be over-dispersed after taking into account the extrinsic noise.

In summary, assuming genes *a* and *b* are intrinsically uncorrelated and the superscript *P ois* denotes intrinsically Poisson distributed genes, these two observations can be expressed as follows:

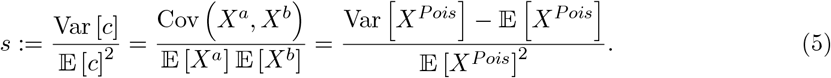

This moment relationship can be validated by estimating extrinsic noise and testing Poisson distributed genes (Section 4.2). Notably, the Poisson distribution property is particularly useful for cell size estimation, as the maximum likelihood estimate (MLE) of the cell size for Poisson genes is simply their sum (Section 4.3). In practice, we filter genes based on a mean expression threshold (>0.1), as the normalized (co)variance of low-expression counts tends to be noisy. We then calculate the normalized covariance between the filtered genes to determine the mean and mode (see Sec 4.2). To determine whether the normalized variance equals the extrinsic noise (average normalized variance), we calculate its bootstrap confidence intervals and the equality holds for a gene if its 95% confidence interval contains the estimated value of extrinsic noise (Sec 4.2). We call those genes “Poisson”.

In the above derivation, we assume that *c*^*bio*^ and *c*^*tech*^ are the same for all genes or species within a single cell. However, this may not be the case and we need to validate it on data. Nevertheless, the model can be extended to cases where multiple *c*^*bio*^ and *c*^*tech*^ values exist for different groups of species and genes. In such cases, we simply need to introduce different *c*^*bio*^ and *c*^*tech*^ for each group and all equations still hold.

### 2.2 Validating the technical noise model with pooled RNA

We first validated our technical noise model and the covariance-variance relationships (Eq 3) on scRNA-seq data of pooled RNA from K562 cells with ERCC using inDrop [22]. As the pooled RNA was homogeneous, the in vivo count for each gene in every droplet followed the same Poisson distribution and was mutually independent (Fig 2a). Therefore, there was no biological extrinsic stochasticity, only technical extrinsic noise. The mode of the normalized covariance was expected to be close to the mean and all genes were expected to be Poisson.

**Figure 2.**
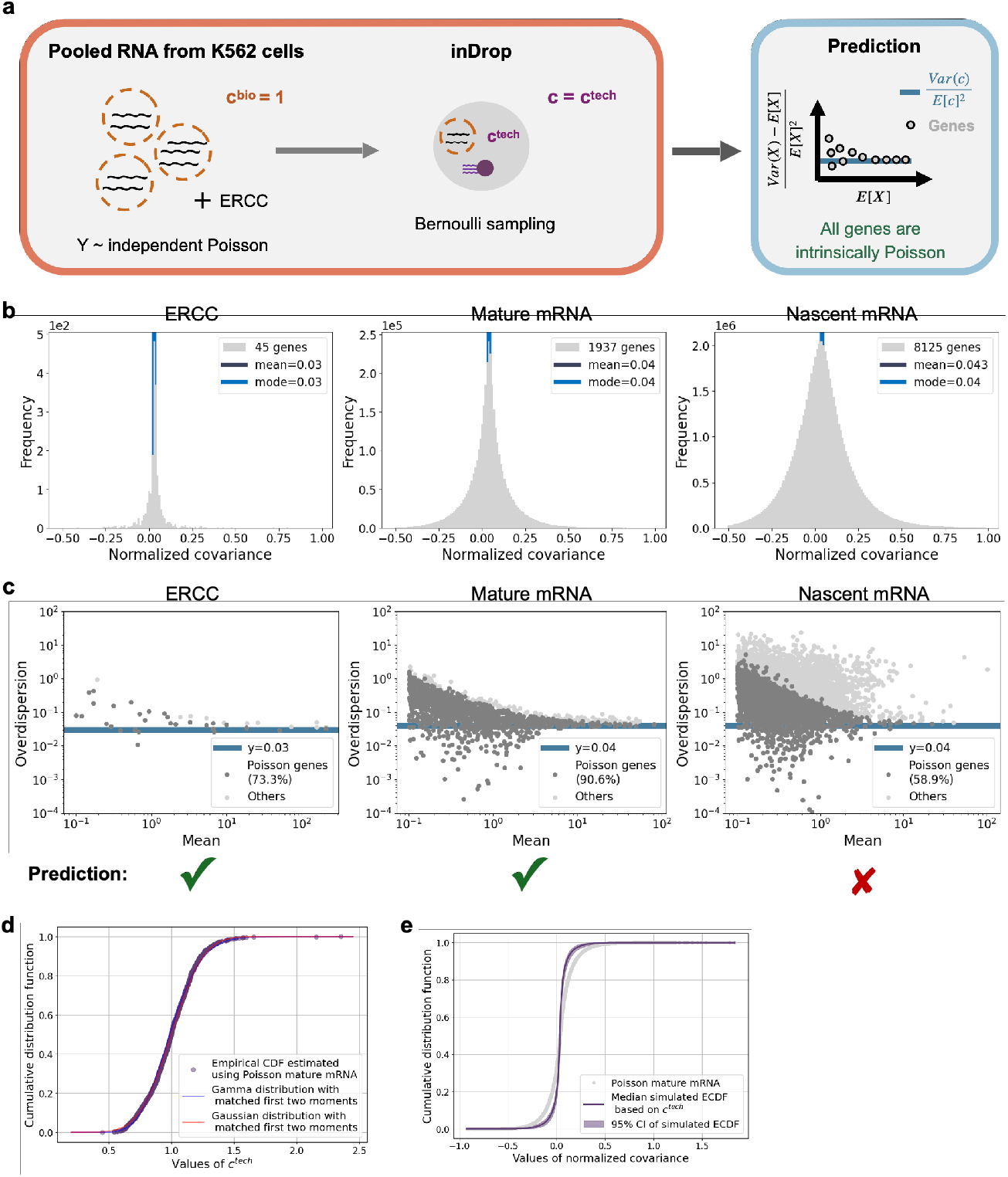
Results for pooled RNA from K562 cells. **a)** Schematic of the experiment and model predictions. **b)** Distribution of normalized covariance between gene pairs with mean expression greater than 0.1, shown separately for ERCC, mature mRNA, and nascent mRNA counts. **c)** Overdispersion-mean relationship for genes with mean expression greater than 0.1, for ERCC, mature mRNA, and nascent mRNA counts, respectively. **d)** Cumulative distribution function of *c*^tech^. Gray dots represent the empirical CDF of estimated *c*^tech^ using selected Poisson mature mRNA counts. The blue line shows a Gamma distribution fitted by matching the first two moments (mean and variance), and the red line shows a Gaussian distribution with the same mean and variance. **e)** Cumulative distribution function of normalized covariance. Gray dots represent the empirical CDF of normalized covariance for mature mRNA counts shown in **b).** Using the estimated *c*^tech^ values and the mean expression levels of the selected genes, 1000 bootstrap samples were generated. The purple line indicates the median empirical CDF across bootstrap replicates, and the light purple band represents the 95% confidence interval.

We calculated the distribution of normalized covariance within ERCC, mature mRNA and nascent mRNA respectively to see if they shared the same *c*^*tech*^ and extrinsic noise (Fig S1a). We found that ERCC and endogenous mRNA seemed to have slightly different *c*^*tech*^, as the estimated extrinsic noise value of ERCC was slightly smaller that that of mRNA. Within the endogenous mRNA, the extrinsic noise appeared to be the same. This suggests that the capture mechanism of ERCC in scRNA-seq might be different from endogenous mRNA, though the difference is small.

Next, we tested whether the covariance-variance relationships held for the ERCC, mature mRNA and nascent mRNA respectively. Given the genes are independent, the covariance-variance relationship holds if the Bernoulli sampling model holds. We plotted the normalized variance against mean, and colored Poisson genes on which the relationships held among all genes (Fig 2b). We also calculated the normalized covariance between selected Poisson genes and found that they were consistent (Fig S1b). The Bernoulli sampling model seemed to work well for the ERCC and mature mRNA but not for the nascent mRNA: the normalized variance of almost half of the nascent counts was much noisier than predicted, while most of the mature mRNA (90%) was within the 95% confidence intervals (Fig 2b). This suggests that the nascent mRNA requires a different measurement model than Bernoulli sampling.

We then sought to characterize *c* (= *c*^*tech*^), which represents the cell size factors commonly used in scRNA-seq analysis [1]. We estimated *c*^*tech*^ as the sum of Poisson-distributed mature RNA counts and found that it correlated well with the total count sum, which was expected since there were no differentially expressed genes (Fig S1c). As the negative binomial distribution is commonly used to model mRNA counts in both pseudo and real cells, which implies that cell size follows a gamma distribution given a Poisson distribution of *in vivo* counts, we asked whether *c*^*tech*^ indeed followed a gamma distribution. We plotted and compared the empirical cumulative distribution function (CDF) of estimated *c*^*tech*^, computed from the sum of Poisson-distributed mature RNA counts, with the CDFs of gamma and Gaussian distributions that shared the same first two moments as *c*^*tech*^. We found that the distribution of *c*^*tech*^ followed a gamma distribution reasonably well, but also fit a Gaussian distribution equally well, if not better (Fig. 2d), which is consistent with the fact that the gamma distribution approaches the Gaussian distribution when the shape parameter is large.

To assess whether a single *c*^*tech*^ was shared across all genes, we compared the empirical CDF of the normalized covariance of mature mRNAs with simulations generated from a Poisson distribution coupled with the empirical distribution of *c*^*tech*^ (Fig 2e). The empirical CDF was less sharp than the empirical CDF of simulations, which suggested that a single *c*^*tech*^ could not fully explain the variability, even for mature mRNA. The single *c*^*tech*^ per cell is the average of the distribution of capture rates for different mature mRNAs in a cell. The capture rates did not seem to relate to the expression levels (Fig S1d). Nevertheless, based on the consistency between the mean and mode of the normalized covariance between all genes and the selected Poisson genes, we concluded that a single *c*^*tech*^ while not entirely accurate, provided a useful approximation.

In summary, homogeneous pooled RNA data revealed that ERCCs, mature mRNAs, and nascent mRNAs are captured through distinct mechanisms, which leads to varying levels of extrinsic noise within ERCCs and endogenous mRNAs and a significantly higher variance in nascent mRNA than would be expected under a simple Bernoulli sampling model. We therefore advocate for more control experiments of this kind to validate technical noise models prior to large-scale data generation.

### 2.3 Validating the “biological” extrinsic noise of pooled RNA mixture

To validate our interpretation of extrinsic noise using a heterogeneous pooled RNA we examined CEL-seq2 data that contained ERCC spike-ins alongside varying amounts of endogenous RNA [23]. The experiment comprised eight distinct RNA mixtures extracted from three human lung cancer cell lines, each present at four RNA amounts within different wells (Fig 3a). Here, the eight RNA mixtures represented eight cell types and the four RNA amounts. Since cell counts were derived from wells containing RNA in solution, we assumed that they were independent and Poisson distributed. However, the different RNA amounts could give rise to biological extrinsic noise and the cell type specific mean parameters could lead to intrinsic variance and covariance. Assuming that the three human lung cancer cell lines have similar concentrations for most genes, we expected the biological extrinsic noise to arise mostly from the variation of RNA amounts. Specifically, the biological extrinsic noise 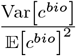 equals the CV^2^ of RNA amounts, which could be calculated to be approximately or slightly above 0.33 based on the experimental design (Section 4.4). Furthermore, those genes that did not vary across the three human lung cancer cell lines were intrinsically Poisson, meaning they had a normalized variance equal to the extrinsic noise, similar to a homogeneous pooled RNA. In contrast, genes that did vary displayed greater dispersion than intrinsically Poisson genes, resulting in a normalized variance that exceeded the extrinsic noise (Prediction in Fig 3a).

**Figure 3.**
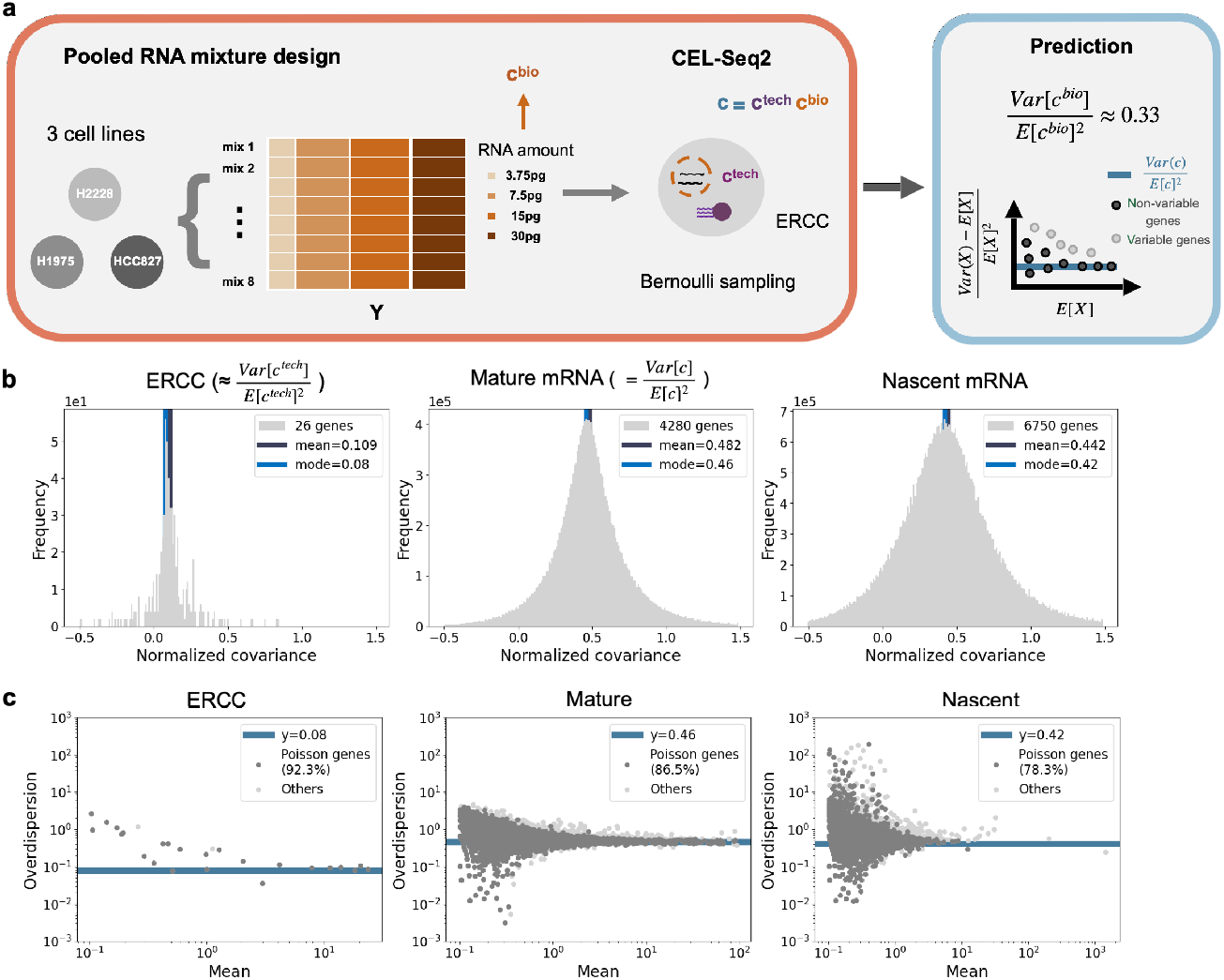
Results for pooled RNA mixture. **a)** Schematic of the experiment and model predictions. **b)** Distribution of normalized covariance between gene pairs with mean expression greater than 0.1, shown separately for ERCC, mature mRNA, and nascent mRNA counts. **c)** Overdispersion-mean relationship for genes with mean expression greater than 0.1, for ERCC, mature mRNA, and nascent mRNA counts, respectively.

We estimated the technical extrinsic noise using the ERCC spike-ins, and also estimate the total extrinsic noise using mature mRNA and nascent mRNA respectively, to see if they were the same. The total extrinsic noise differed within mature and nascent mRNA: the estimated extrinsic noise was slightly higher for mature mRNA (0.46) than nascent mRNA (0.42) (Fig 3b). The distribution of normalized covariance between mature and nascent mRNA had a similar mode to that of nascent mRNA (Fig S2a), indicating that the extrinsic noise of mature mRNA included both components shared with nascent mRNA and components unique to mature mRNA. Most ERCC (92%) and mature mRNA (87%) fell within the 95% confidence intervals and satisfied the Poisson criteria, whereas nascent mRNA counts had a smaller percentage (78%) and were noisier with larger normalized variance (Fig 3b). As a consistency check, the normalized covariance between Poisson genes showed similar values, with differences within 0.01 (Fig S2b). Based on these observations, we speculate that the capture of mature and nascent mRNA shared similar mechanisms as well as distinct differences, which led to the small difference in extrinsic noise. Importantly, nascent mRNA is likely to require a slightly nosier technical model than Bernoulli sampling.

Therefore, we used normalized covariance of mature mRNA for calculating the total extrinsic noise and ERCC for the technical extrinsic noise to estimate the biological extrinsic noise based on equation 4. The estimated biological extrinsic noise was 0.35, which was reasonably closed to the expectation (0.33). Then we calculated the total and technical cell size (*c*_*i*_ and 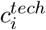) using the sums of Poisson mature mRNA and ERCC respectively, which were similar to those using total counts (Fig S2c). We estimated biological cell size 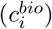 by taking the ratio of *c*_*i*_ and 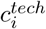, and compared the distribution of 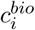 to the expected distribution from the experimental design (Section 4.4). Cells were centered around the expected values but the variance seemed to be large (Fig S2d and e).

### 2.4 Decomposing biological and technical extrinsic noise using species-mixing experiments

Based on the results obtained with RNA in solution, we decided to focus on the mature counts. However, in this context of experiments with individual cells, the logic is reversed. Unlike in pooled RNA, where genes can be assumed to follow a Poisson distribution in pseudocells, these assumptions do not inherently hold *in vivo*. Instead, by testing the relationship between extrinsic noise and overdispersion across genes, we identified those genes for which the assumptions of the Poisson distribution hold, at least approximately. This enables genome-scale understanding of gene expression noise and provided an additional piece of evidence in the context of previously inconsistent findings regarding biological variability [24, 25]

Therefore, we used an iterative approach to estimate extrinsic noise and to identify “Poisson” genes (Fig S3). Starting with all genes, we calculated the normalized covariance and estimated the extrinsic noise, which was then used to identify genes whose normalized variances were close to the extrinsic noise. Next, we re-estimated the extrinsic noise using the normalized covariance between selected genes and compared it to the previous value. This process was repeated iteratively until the extrinsic noise estimate stabilized (with differences within 10%), though typically, at most one iteration was needed. Therefore, we assumed that these selected genes were independent (on average) and had intrinsic variance similar to a Poisson distribution, so we referred to them as “Poisson”. Although their exact distributions may deviate from a Poisson model, these genes were likely to exhibit low variability across cells, rendering them appropriate for estimating cell size.

We first sought to characterize the biological and technical contribution of extrinsic noise. To measure technical extrinsic noise, we needed some control RNA in the same cell. Usually, ERCC spike-ins are used as external controls to quantify technical variance. However, here we utilized the ambient mRNA as the control mRNA by leveraging species-mixing experiments, which are commonly used to assess doublet rates. In these scRNA-seq experiments, human and mouse cells are typically mixed, resulting in ambient mRNA from both species in droplets containing cells from only one species. Then the ambient mRNA from the other species serves as an external RNA control for technical extrinsic noise (Fig. 4a).

**Figure 4.**
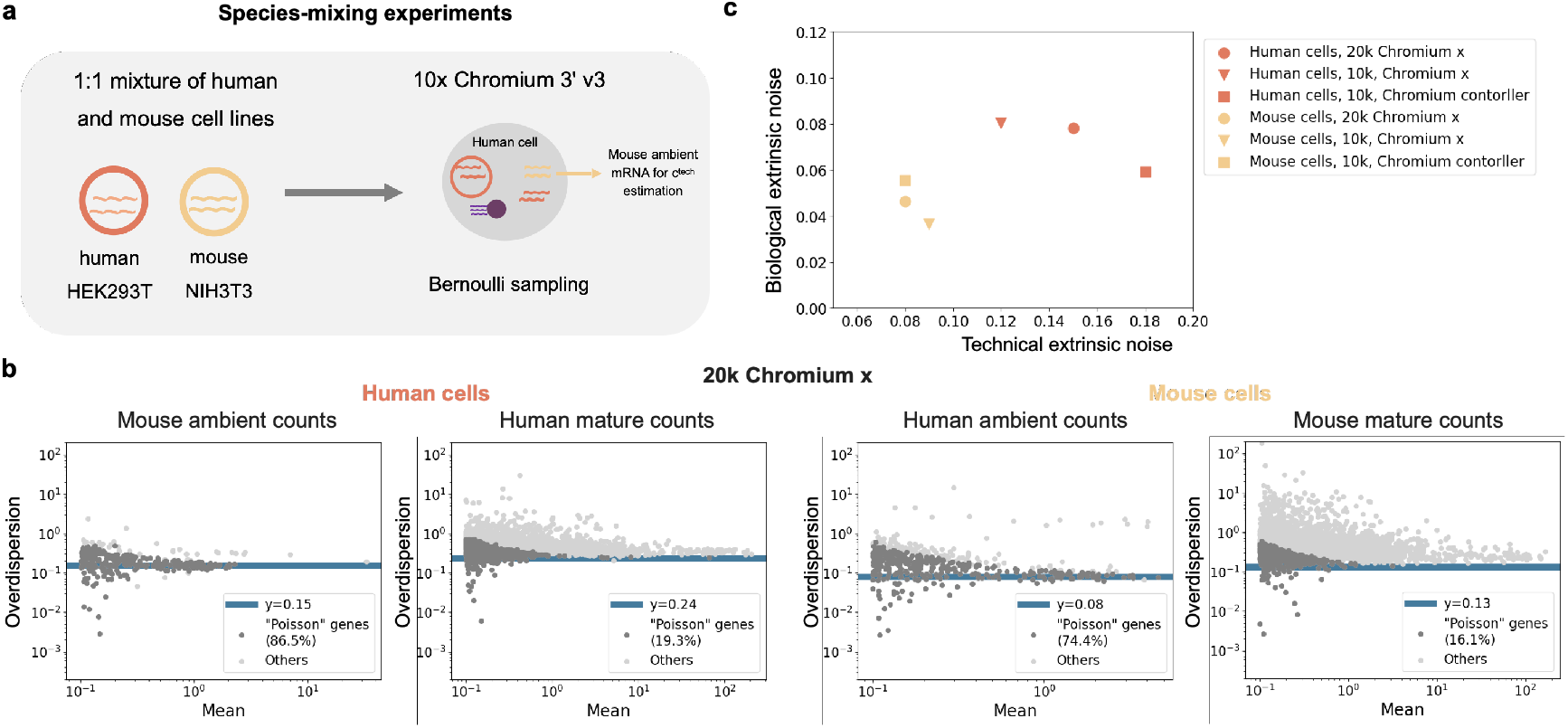
Extrinsic noise in species-mixing experiments. **a)** Schematic of the species-mixing experiment. **b)** Overdispersion-mean relationships for human and mouse genes in both human and mouse cells in the 20k Chromium X dataset. **c)** Biological and technical extrinsic noise in three species-mixing experiments.

In light of this, we calculated the biological and technical extrinsic noise of three 10x human-mouse mixture datasets. We used mature mRNA of the corresponding species to estimate total extrinsic noise, and total ambient mRNA from the other species to estimate technical extrinsic noise. We did not distinguish between nascent and mature and used the total counts when calculating normalized covariance because their counts are low. For example, in droplets containing only human cells, the technical extrinsic noise estimated from mouse ambient mRNA is 0.15, and, similar to pooled RNA, the genes were expected to follow a Poisson distribution. In contrast, the total extrinsic noise estimated from human mRNA is 0.24, with fewer than 20% of human transcripts falling within the Poisson range (Fig 4b). The differing percentages between ambient and cellular mRNA highlight that only a small fraction of in vivo transcript counts potentially follow a Poisson distribution. Based on the total and technical extrinsic noise, we estimate the biological extrinsic noise based on equation 4, which leads to 0.15 for human cells (Fig 4b). Both the biological and technical contributions to extrinsic noise are substantial (Fig 4c). The estimated biological extrinsic noise are relatively robust across three datasets, and the mouse cells (NIH3T3) seems to be slightly more homogeneous than human cells (HEK293T).

### 2.5 Characterizing extrinsic noise and cell size factors on scRNA-seq data

Given the non-negligible contribution of biological extrinsic noise even in homogeneous cell lines, we argue that even when biological extrinsic noise cannot be explicitly distinguished due to the absence of control mRNA, a substantial portion of the total extrinsic noise likely reflects underlying biological variation and should not be disregarded. Therefore, we applied our procedure to several scRNA-seq datasets and characterized both extrinsic noise and cell size factors.

We first investigated whether extrinsic noise is related to the average abundance of genes, a topic that has been debated in previous studies [8, 26]. For our analysis, we selected the 10x Flex K562 dataset because the probes covering exon junctions yield more abundant mature mRNA counts. We then plotted the distribution of normalized covariance across genes with varying mean expression levels and found that the modes of the distributions were identical (0.23), with comparable means (Fig. 5a). The percentage of “Poisson” genes is also comparable to those observed in the species-mixing data (Fig. 5b). The distributions of normalized covariance across “Poisson” genes with varying mean expression levels also show no difference with the same modes (Fig. S5a). We repeated the same analysis on STORM-seq K562 dataset which also has abundant mature counts [27], and find that extrinsic noise (the mode of normalized covariance) is also the same (0.24) across genes with different abundance (Fig. S6a). We concluded that extrinsic noise and the resulting baseline overdispersion are not related to the average abundance of genes.

**Figure 5.**
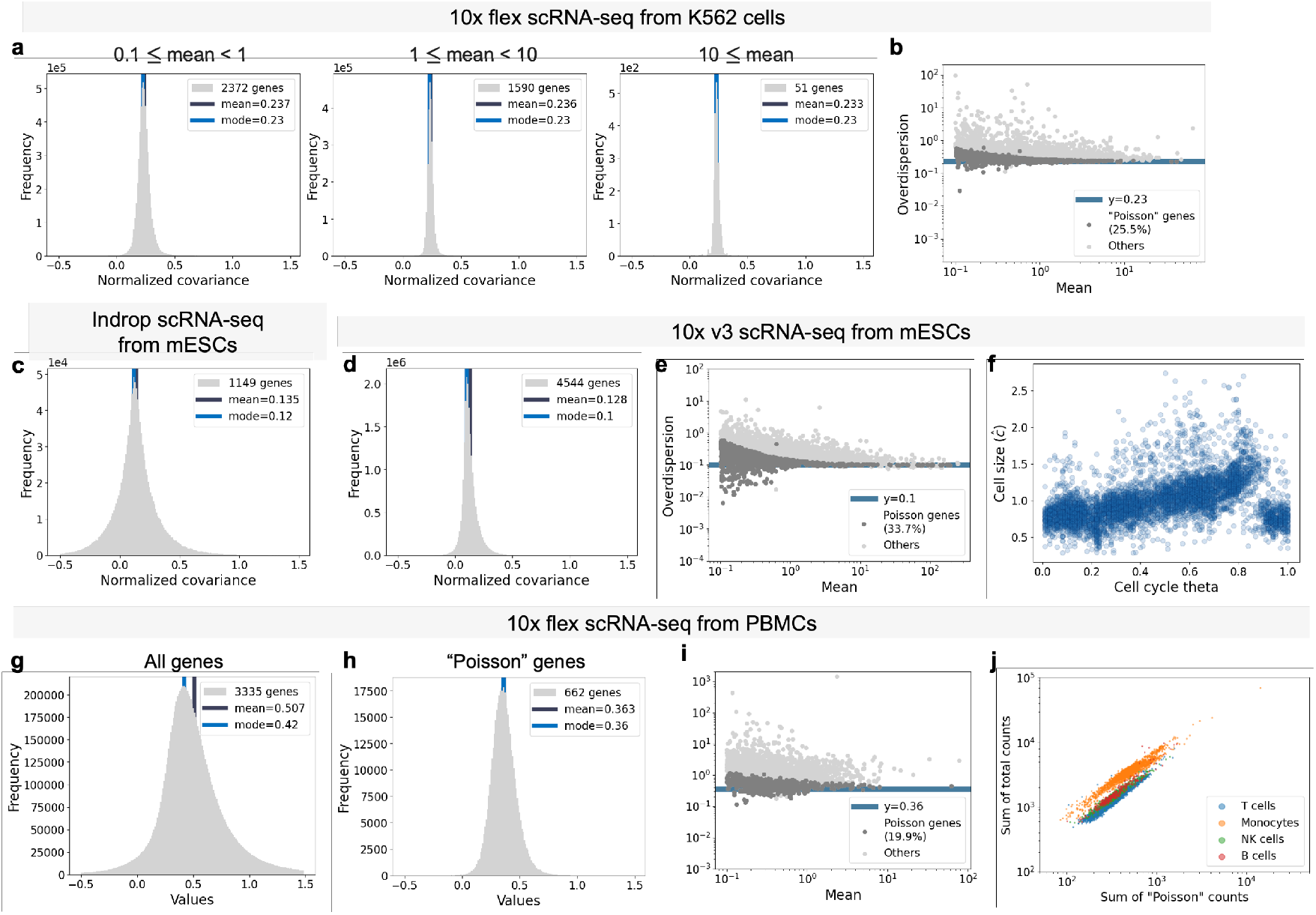
Extrinsic noise in single cell datasets. **(a)** Distribution of normalized covariance within genes across different mean expression ranges for the K562 10x Flex dataset. **(b)** Overdispersion-mean relationship for the K562 10x Flex dataset. **(c)** Distribution of normalized covariance for the mESC inDrop dataset. **(d)** Distribution of normalized covariance for the mESC 10x 3’ v3 dataset. **(e)** Overdispersion-mean relationship for the mESC 10x 3’ v3 dataset. **(f)** Cell size along cell cycle. Cell sizes are estimated using Poisson genes from panel (e). Cell cycle progression is denoted by cell cycle theta, as reported by Riba et al. (2022). **(g)** Distribution of normalized covariance between gene pairs with mean expression greater than 0.1 for the PBMC dataset. **(h)** Distribution of normalized covariance between selected Poisson genes for the PBMC dataset. **(i)** Overdispersion-mean relationship for the PBMC dataset. **(i)** Sum of total counts against sum of Poisson counts, colored by cell types.

We then investigated whether extrinsic noise is associated with cell cycle progression. To do this, we used mouse embryonic stem cells (mESC) data with inferred cell cycle stages [28]. We estimated the extrinsic noise (Fig. 5c) and also compared it to that of mESC inDrop data [22], finding that the estimates were similar (Fig. 5c). This indicates the robustness of the extrinsic noise across different datasets. We selected Poisson genes (Fig 5e), and estimate *ĉ* as the sum of “Poisson” mature RNA counts, which correlates well with the total count sum (Fig S7c). We plotted the estimated cell size factors (*ĉ*) along the inferred transcriptional phase (cell cycle *θ*) from Riba et al. and observed a clear pattern of cell size variation across the cell cycle (Fig. 5f), which confirms that the cell cycle contributes to extrinsic noise.

Up to this point, the sum of Poisson counts had shown a perfect correlation with the total counts. However, this may not be the case for heterogeneous cells, i.e., datasets consisting of different cell types. To investigate this, we applied our approach to the peripheral blood mononuclear cells (PBMC) dataset generated using 10x flex technology [29]. We found that the extrinsic noise in PBMCs is significantly higher than that observed in homogeneous cell types such as mESC and K562. The distribution of normalized covariance across all genes is right-skewed (Fig. 5g), suggesting that many genes exhibit positive correlations. On the other hand, the distribution across Poisson genes is more symmetrical, with the mode and mean closely aligned (Fig. 5h). The percentage of “Poisson” genes remains similar (Fig 5i). However, the estimated *ĉ* as the sum of “Poisson” mature RNA counts no longer aligns well with the total count sum (Fig 5j). We used the Leiden algorithm to cluster cells into T cells, Natural killer (NK) cells, B cells, and Monocytes based on marker genes. Different cell types within PBMC appear to have varying ratios of “Poisson” to total counts (Fig 5j), likely reflecting the presence of highly differentially expressed genes specific to monocytes. Therefore, using “Poisson” counts or total counts will result in different cell size factors. To demonstrate the impact on downstream analysis, we normalized the raw counts using both the total UMI count and the sum of Poisson gene counts, respectively, and performed a Mann–Whitney U test to identify differentially expressed genes between monocytes and natural killer cells. We compared the resulting p-values (Fig. S8a), and listed the top 100 differentially expressed genes identified under each normalization method for comparison (Fig. S8b). We found that this phenomenon is dataset-specific and depends on the underlying cellular composition, as demonstrated by the 10x mouse forebrain data [30], where the sum of ‘Poisson’ mature RNA counts aligns better with the total count sum (Fig. S9d).

## 3 Discussion

In this work, we clarify the underlying assumptions of extrinsic noise in scRNA-seq normalization and describe an extrinsic noise model that has only been implicitly recognized in previous studies. This model establishes a direct relationship among normalized covariance, extrinsic noise, and the overdispersion observed in intrinsically Poisson genes following previous study [14]. This relationship enables us to validate the model using pooled RNA data and to identify genes whose expression variance is consistent with a Poisson distribution. By providing a baseline for overdispersion, extrinsic noise reveals that much of the observed overdispersion in scRNA-seq data can still be explained by genes that are intrinsically Poisson.

We have shown how a mechanistic model can lead to testable predictions, and how validating these predictions can either support the model or prompt the development of alternative explanations [31]. Specifically, we found that Bernoulli sampling is applicable only to mature RNA counts, likely due to differences in capture mechanisms between mature and nascent mRNA. Even for mature counts, using a single cell size factor remains a coarse approximation. Furthermore, given that the technical noise model may vary across species and technologies, we advocate for more careful assessment in future experimental designs, recommending that the technical noise model be characterized prior to large-scale data generation.

Beyond quantifying biological and technical extrinsic noise, a key motivation for modeling extrinsic noise and cell size is to enhance the accuracy of downstream data analysis. Rather than simply normalizing total counts by cell size, we advocate for explicitly incorporating the cell size factor when modeling variable genes with more complex gene expression models, such as those beyond constitutive expression and Poisson distributions. The cell size estimators derived from Poisson genes can be treated as constants and provided as inputs to the inference process, thereby simplifying the modeling of other variable genes. Because these estimators are based on two orthogonal groups of genes— those used for cell size estimation and those being modeled-this approach effectively avoids the issue of “double-dipping” and ensures a more robust and reliable analysis.

We have provided only preliminary insights into extrinsic noise, as our analysis is based on modeling single genes and does not specify a detailed gene expression model. As a result, we cannot determine the exact forms of variance and covariance beyond what is expected from a Poisson distribution. For genes that follow Poisson statistics, all biological extrinsic noise arises from variation in their mean expression levels. In such cases, it is sufficient to decompose extrinsic noise into biological and technical components. However, for more variable genes that exhibit super-Poissonian variance, more sophisticated models such as bursty transcription are needed to accurately capture their expression dynamics [32]. These models introduce additional parameters, such as burst frequency and burst size, to account for the excess variability. While most studies assume that only burst size scales with cell size [14, 20, 21], we show that biological extrinsic noise can, in fact, be further decomposed. This allows for a more detailed, quantitative dissection of how each parameter contributes to extrinsic noise (see Section 4.5). We advocate for future studies to adopt more comprehensive modeling approaches in order to deepen our understanding of the sources and mechanisms underlying gene expression variability.

## 4 Methods

### 4.1 Extrinsic noise model

Extrinsic noise is global to a cell and contains both biological and technical components. Therefore, we introduce two random variables to represent biological and technical cell factors. Specifically, for the biological model, we assume that each cell has a random variable *c*^*bio*^ and the mean of every gene is proportional to this value. *c*_*bio*_ could result from the cell volume, cell cycle and factors that effect all genes. Without loss of generality, we assume 𝔼 [*c*_*bio*_] = 1. Denote the in vivo number of gene j in cell i by 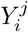, this model for biological extrinsic noise means

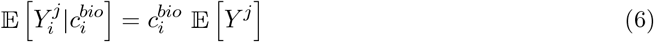

where 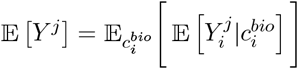 is the mean of gene j across cells.

As consequence of law of total variance, for covariance and variance of gene a and b across cells, we have

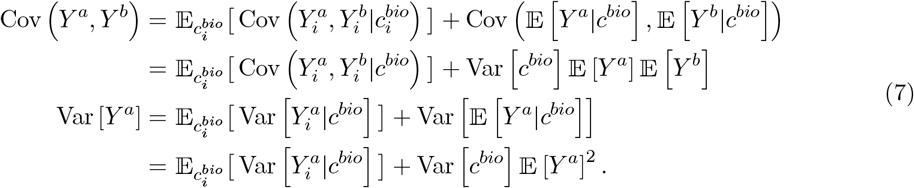

The first terms on the right-hand side of both equations describe the intrinsic covariance and variance, which depend on and also reflect the gene expression mechanism. The second terms describe the extrinsic noise introduced by *c*^*bio*^. This separation of intrinsic and extrinsic terms has been addressed in previous studies [9, 11, 12].

For a technical noise model, we assume Bernoulli sampling of transcripts in single-cell sequencing experiments, which leads to binomial distribution of observed counts given in vivo counts. Similarly we assume each cell has a random variable *c*_*tech*_ and the success probability in binomial distribution is proportional to this value. This *c*_*tech*_ could be interpreted as relative read depth during sequencing and is independent of the biological model and in vivo counts. We introduce a constant capture rate *λ* for each species of molecule so that we can again assume 𝔼 [*c*_*tech*_] = 1. Denote the observed counts after single cell sequencing by *X*, this technical model means

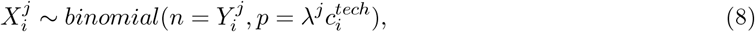

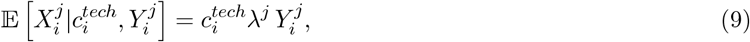

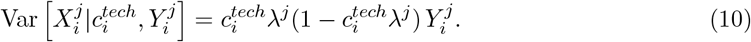

Using law of total variance (covariance) gives

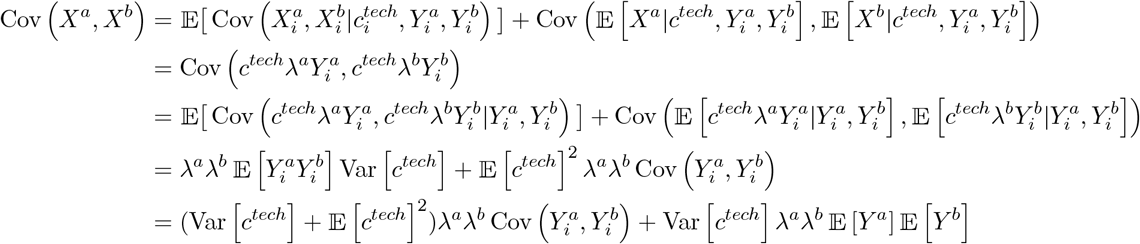

and

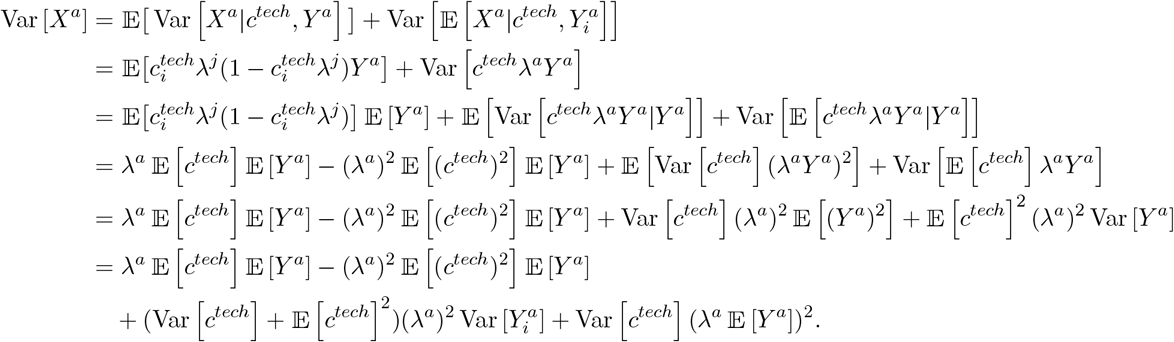

Therefore,

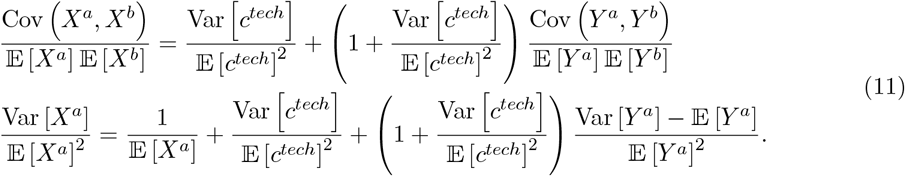

Plugging in equation 7, we arrive at the expression that connects intrinsic and observed noise under our extrinsic noise model,

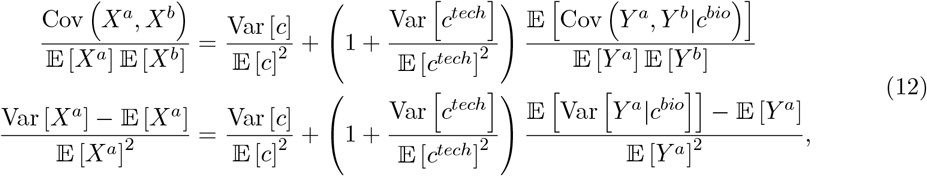

where *c* = *c*^*tech*^*c*^*bio*^ is the overall cell factor. This expression is rather general and follows directly from equation 1 and equation 2. The shared constant factors 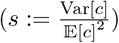 denote the extrinsic noise, and potentially explain the constant offset observed in the plot of the Fano factor against mean for genes. We denote 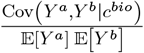 and 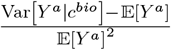 by intrinsic covariance and variance.

### 4.2 Procedure for estimating extrinsic noise and selecting Poisson genes

To estimate the extrinsic noise, we calculated the normalized covariance between genes with mean expression greater than 0.1, and used the mode of the resulting distribution. This was computed using histogram bins of width 0.01. The center of the bin with the highest frequency was taken as the estimated value, which was set to exactly two decimal digits by construction of the bin edges.

To select Poisson genes, we calculated the 95% bootstrap interval of overdispersion for each gene based on 1000 bootstrap samples by default. Then we selected genes whose 95% bootstrap intervals contain the estimated extrinsic noise.

### 4.3 Maximum likelihood estimation of cell size

Let 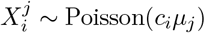 be the observed expression count of gene *j* in cell *i*, where *c*_*i*_ is the cell size (scaling factor) for cell *i* and *µ*_*j*_ is the mean of gene *j*.

The likelihood function for cell *i* given its gene expression vector 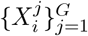 is given by

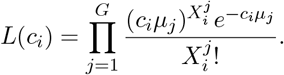

The log-likelihood is given by

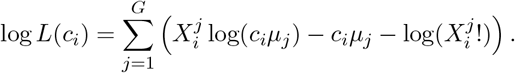

To find the maximum likelihood estimator (MLE) of *c*_*i*_, we differentiated the log-likelihood with respect to *c*_*i*_ and set the derivative to zero:

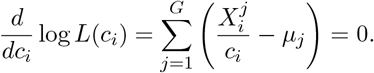

Solving for *c*_*i*_ gives the MLE:

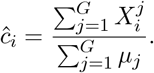

### 4.4 The expected biological extrinsic noise of the CEL-seq2 data

The RNA mixture was prepared on a 384-well plate (Supplementary Figure 1a in [23]). After processing the SRA files from GEO Series GSE117617 using kb-python [33] and filtering out two outlier cells, we obtained a final dataset consisting of 357 cells. The exact biological extrinsic noise is influenced by the RNA amounts of the remaining 357 cells, which remain unknown. However, assuming that the 27 removed cells each had an RNA amount of 3.75 *µ*g, we can estimate a lower bound for the biological extrinsic noise, which is approximately 0.333.

### 4.5 Identifiability of parameter-specific extrinsic noise in bursty models

Here we consider the bursty transcription model, and assume a random variable for the extrinsic noise associated with each parameter. Studying the identifiability of parameter-specific extrinsic noise is crucial, as it can help us understand how biological extrinsic noise influences gene expression. By investigating this aspect, we aim to gain insights into the mechanisms that drive variability in gene expression at the single-cell level.

We consider the following bursty model of nascent and mature mRNA:

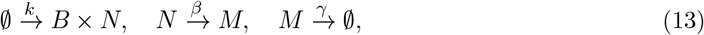

where in the first reaction the number of nascent mRNA molecules synthesized in each burst (B) follows a geometric distribution on {0, 1, 2, …} with a mean of b, referred to as the burst size [34]. The distribution of nascent is well known to be negative binomial, but the joint distribution of nascent and mature counts is not analytically available. Here, following the framework in [35], we use the generating function method to investigate the identifiablity of extrinsic noise, which can be extended to general gene expression models.

Denote the *in vivo* nascent and mature mRNA counts by *y*_*u*_ and *y*_*m*_. Let 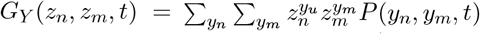 be the generating function. Assuming *β* ≠ *γ*, the factorial generating function *φ*_*Y*_ (*u*_*n*_, *u*_*m*_, ∞) := log *G*_*Y*_ (*u*_*n*_ + 1, *u*_*m*_ + 1, ∞) is [34]:

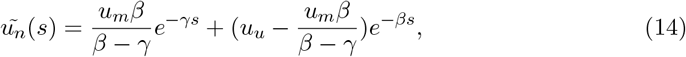

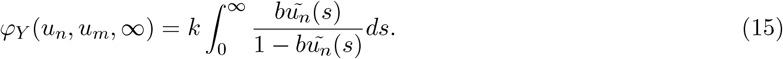

Adding Bernoulli sampling with rate *c*^*tech*^*λ*, where *λ* is the species-specific constant, the generating function of observed counts *x* is

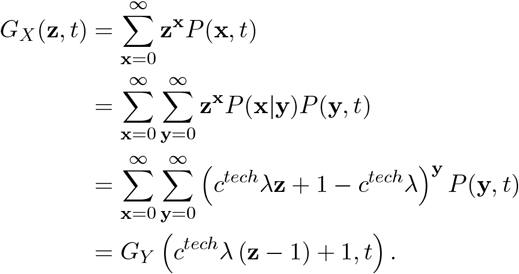

Then, adding parameter-specific extrinsic noise to each parameter, the factorial generating function of observed counts *X φ*_*X*_(*u*_*n*_, *u*_*m*_, ∞) := log *G*_*X*_(*u*_*n*_ + 1, *u*_*m*_ + 1, ∞) is:

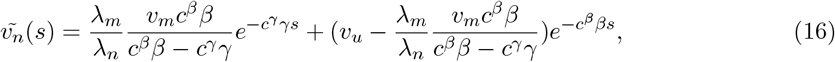

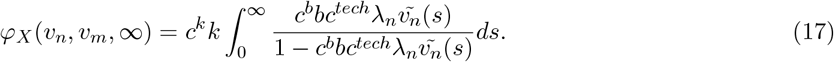

Note that all *c* values are shared across genes within the same cell. Given the identifiable parameters in equation 14 for *in vivo* counts *Y* are *b*, 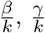 [34], the identifiable parameters in equation 16 from observed counts *X* are *bλ*_*n*_, 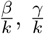, and 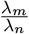, as well as the relative values of 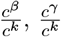, and *c*^*b*^*c*^tech^, under the assumption that

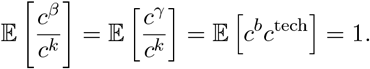

With the use of external RNA controls, it becomes possible to further disentangle *c*^*b*^ from *c*^tech^.

## 5 Data and code availability

All datasets used in this study are publicly available. Raw FASTQ files were downloaded for each dataset and processed using kb-python version 0.29.1 with the nac workflow [36, 33]. The links to FASTQ files are in Supplementary Table S1.

All code used to generate the results and figures in the paper is available at https://github.com/pachterlab/FP_2025.

## 6 Acknowledgments

We thank Delaney Sullivan for help with processing scRNA-seq data. We thank Gennady Gorin and Tara Chari for helpful discussions.

