## Supplementary Information for "Extrinsic biological stochasticity and technical noise normalization of single-cell RNA sequencing data"

#### 1 Supplementary Tables

| Dataset | FASTQs | Reference |
|---|---|---|
| Homogeneous RNA solution (Indrops v1) | GSM1599501 | [1] |
| Heterogeneous RNA solution (CEL-seq2) | GSM3305230 | [2] |
| Species-mixing, 20k Chromium X (10x 3' v3) | `https://s3-us-west-2.amazonaws.com/10x.files/samples/cell-exp/6.1.0/20k_hgmm_3p_HT_nextgem_Chromium_X/20k_hgmm_3p_HT_nextgem_Chromium_X_fastqs.tar` | [3] |
| Species-mixing, 10k Chromium X (10x 3' v3) | `https://s3-us-west-2.amazonaws.com/10x.files/samples/cell-exp/6.1.0/10k_hgmm_3p_nextgem_Chromium_X/10k_hgmm_3p_nextgem_Chromium_X_fastqs.tar` | [4] |
| Species-mixing, 10k Chromium controller (10x 3' v3) | `https://s3-us-west-2.amazonaws.com/10x.files/samples/cell-exp/6.1.0/10k_hgmm_3p_nextgem_Chromium_Controller/10k_hgmm_3p_nextgem_Chromium_Controller_fastqs.tar` | [5] |
| K562 (10x flex) | `https://s3-us-west-2.amazonaws.com/10x.files/samples/cell-exp/7.0.0/10k_K562_singleplex_Multiplex/10k_K562_singleplex_Multiplex_fastqs.tar` | [6] |
| mESC (10x 3' v3) | GSM5111566 | [7] |
| mESC (Indrops v1) | GSM1599494 | [1] |
| PBMC (10x flex) | `https://s3-us-west-2.amazonaws.com/10x.files/samples/cell-exp/8.0.0/10k_Human_PBMC_TotalSeqB_singleplex_Multiplex/10k_Human_PBMC_TotalSeqB_singleplex_Multiplex_fastqs.tar` | [8] |
| Mouse forebrain (10x flex) | `https://cf.10xgenomics.com/samples/cell-exp/7.1.0/10k_mouse_forebrain_scFFPE_singleplex_Multiplex/10k_mouse_forebrain_scFFPE_singleplex_Multiplex_fastqs.tar` | [9] |
| K562 (STORM-seq) | GSE296405 | [10] |

**Table S1: Datasets metadata.** Datasets used for all analyses, with their technology in parentheses, FASTQ files accession links, and references.

### 2 Supplementary Figures

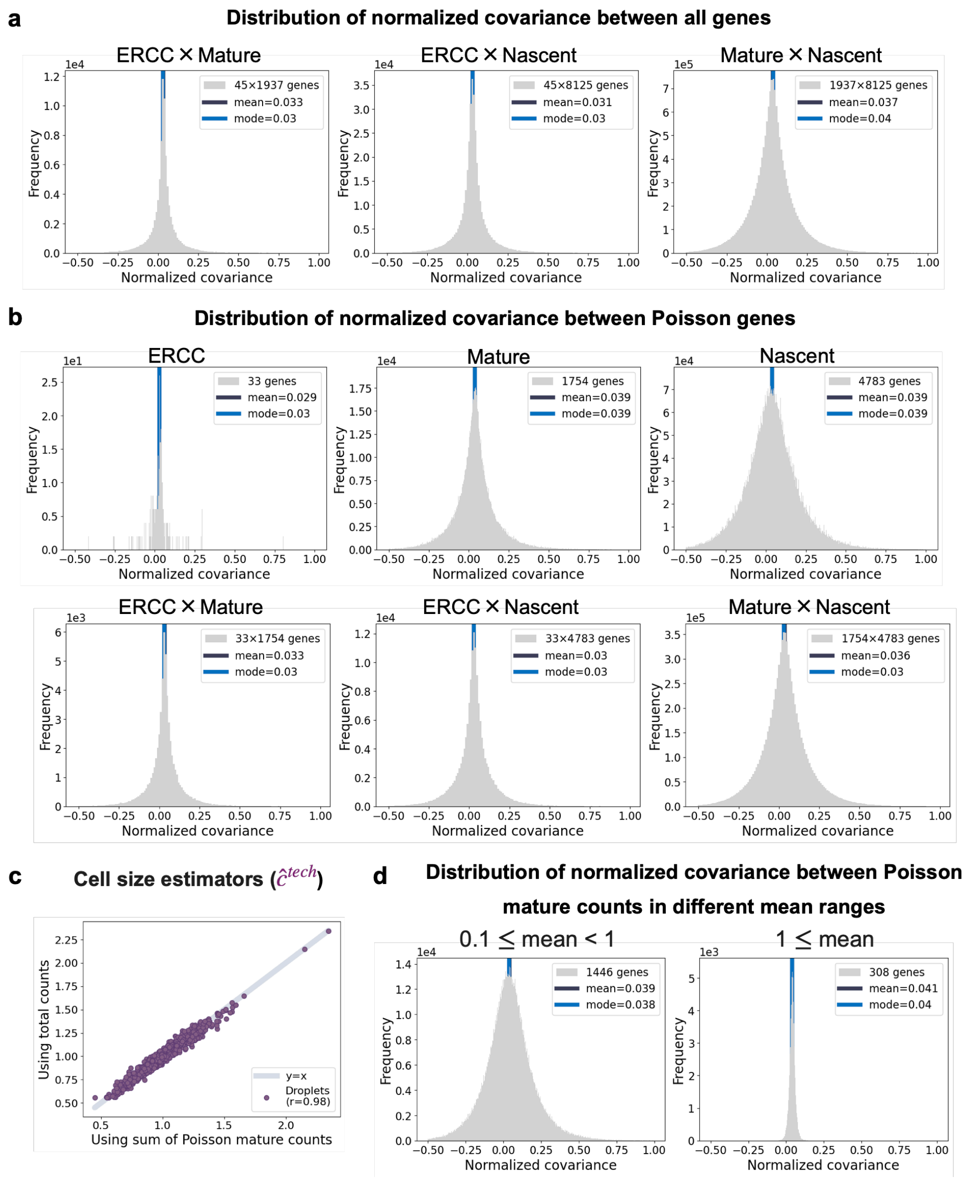


**Figure S1: Supplementary figures for homogeneous RNA solution (a)** Distribution of normalized covariance between different species. **(b)** Distribution of normalized covariance between Poisson genes of different species. **(c)** Comparison of cell size estimators using the sum of total counts and Poisson mature counts. **(d)** Distribution of normalized covariance between Poisson mature counts across different mean expression ranges.

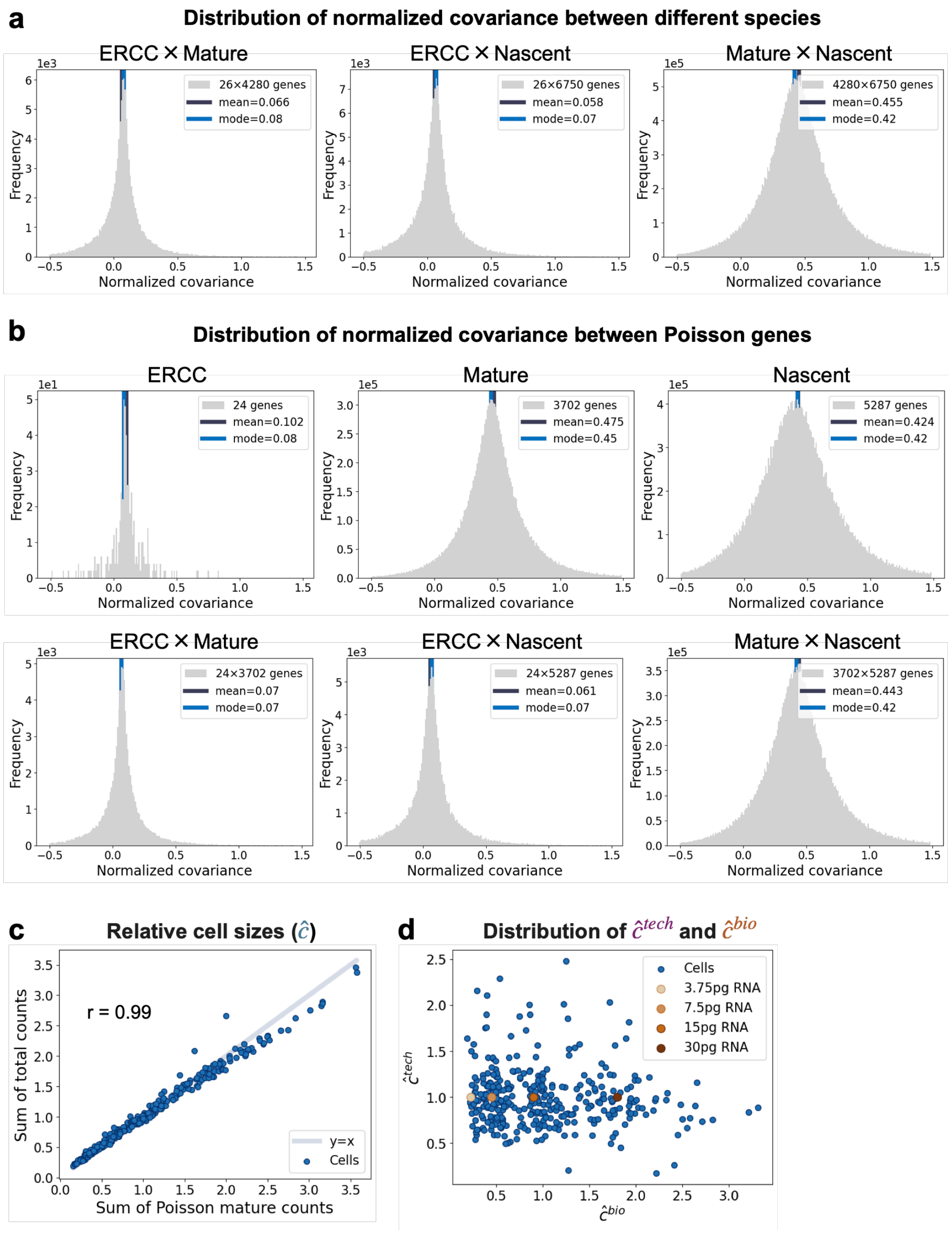


**Figure S2: Supplementary figures for heterogeneous RNA solution (a)** Distribution of normalized covariance between different species. **(b)** Distribution of normalized covariance between Poisson genes of different species. **(c)** Comparison of cell size estimators using the sum of total counts and Poisson mature counts. **(d)** Distribution of $c^{\text{tech}}$ and $c^{\text{bio}}$. The values of $c^{\text{tech}}$ and $c$ are estimated from the total Poisson ERCC counts and mature mRNA counts, respectively. Based on these estimates, $c^{\text{bio}}$ is computed. The brown dots are the theoretical $c^{\text{bio}}$.

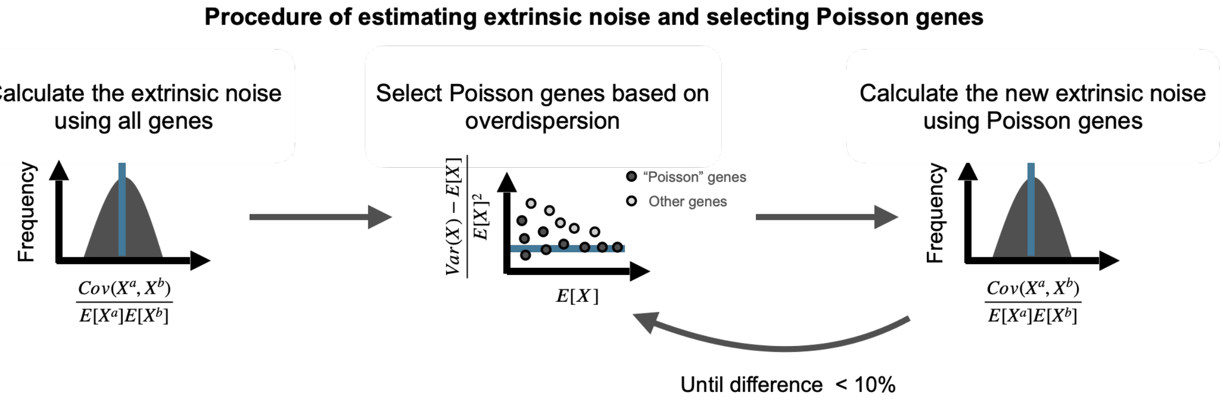


**Figure S3: Procedure on single cell datasets.**

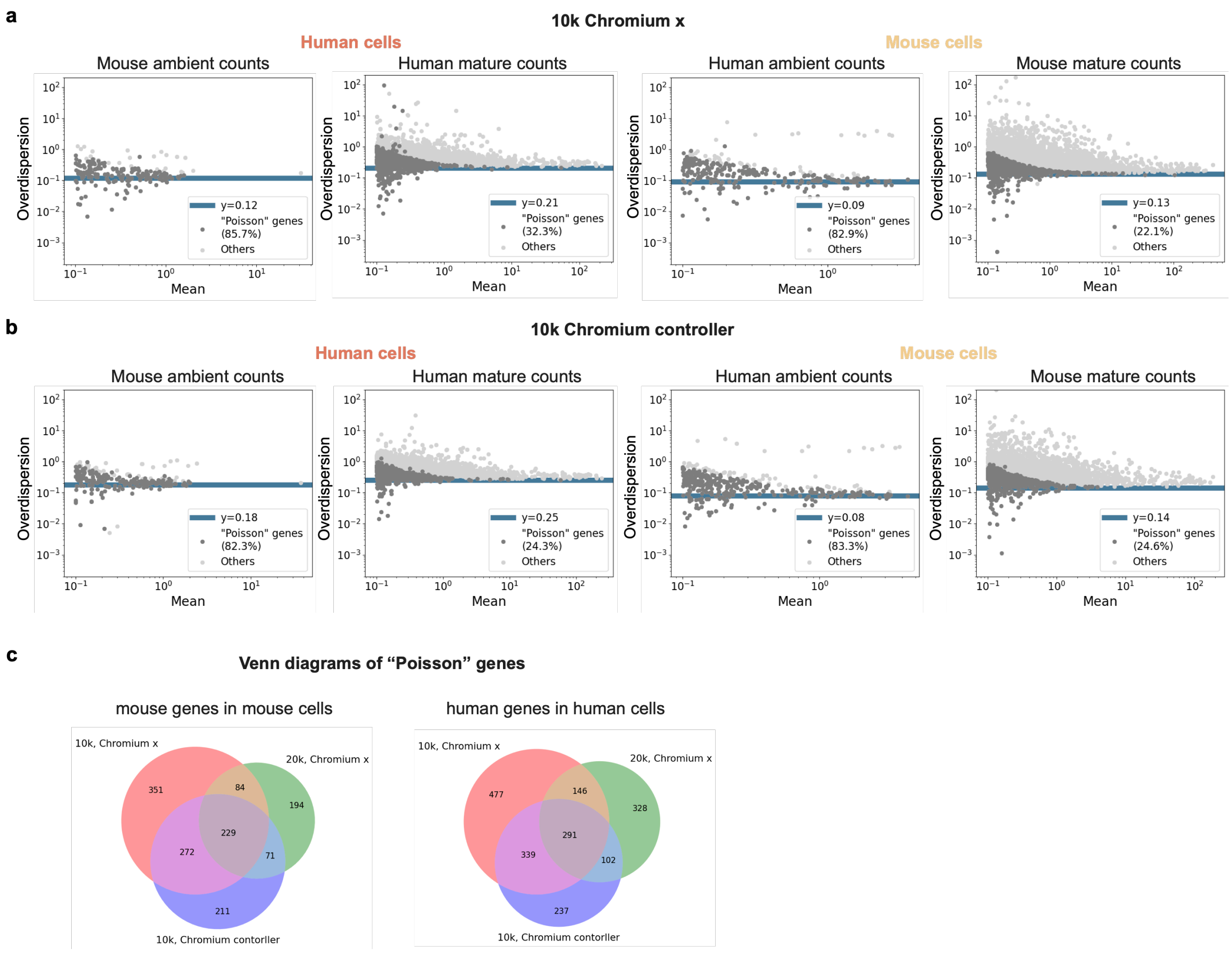


**Figure S4: Supplementary figures for species-mixing experiments (a)** Overdispersion-mean relationships for human and mouse genes in both human and mouse cells in the 10k Chromium X dataset. **(b)** Overdispersion-mean relationships for human and mouse genes in both human and mouse cells in the 10k Chromium controller dataset. **(c)** Venn diagram of selected Poisson genes across the three datasets.

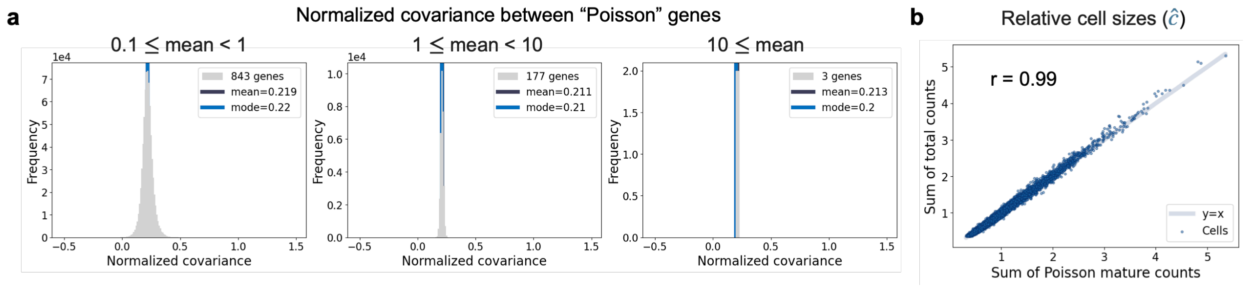


**Figure S5: Supplementary figures for K562 10x flex dataset a)** Distribution of normalized covariance between Poisson mature counts across different mean expression ranges. **b)** Comparison of cell size estimators using the sum of total counts and Poisson mature counts. r denotes the Pearson correlation coefficient.

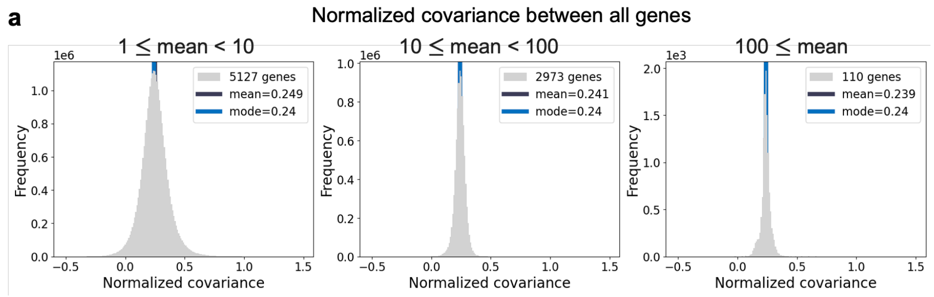


**Figure S6: Results for K562 STORM-seq dataset.(a)** Distribution of normalized covariance between genes across different mean expression ranges.

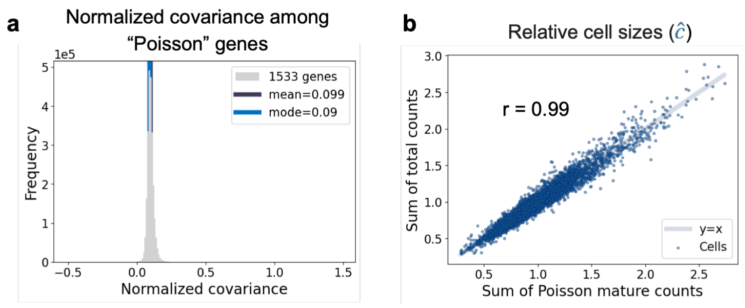


**Figure S7: Supplementary figures for mESC 10x dataset a)** Distribution of normalized covariance between Poisson mature counts. **b)** Comparison of cell size estimators using the sum of total counts and Poisson mature counts. r denotes the Pearson correlation coefficient.

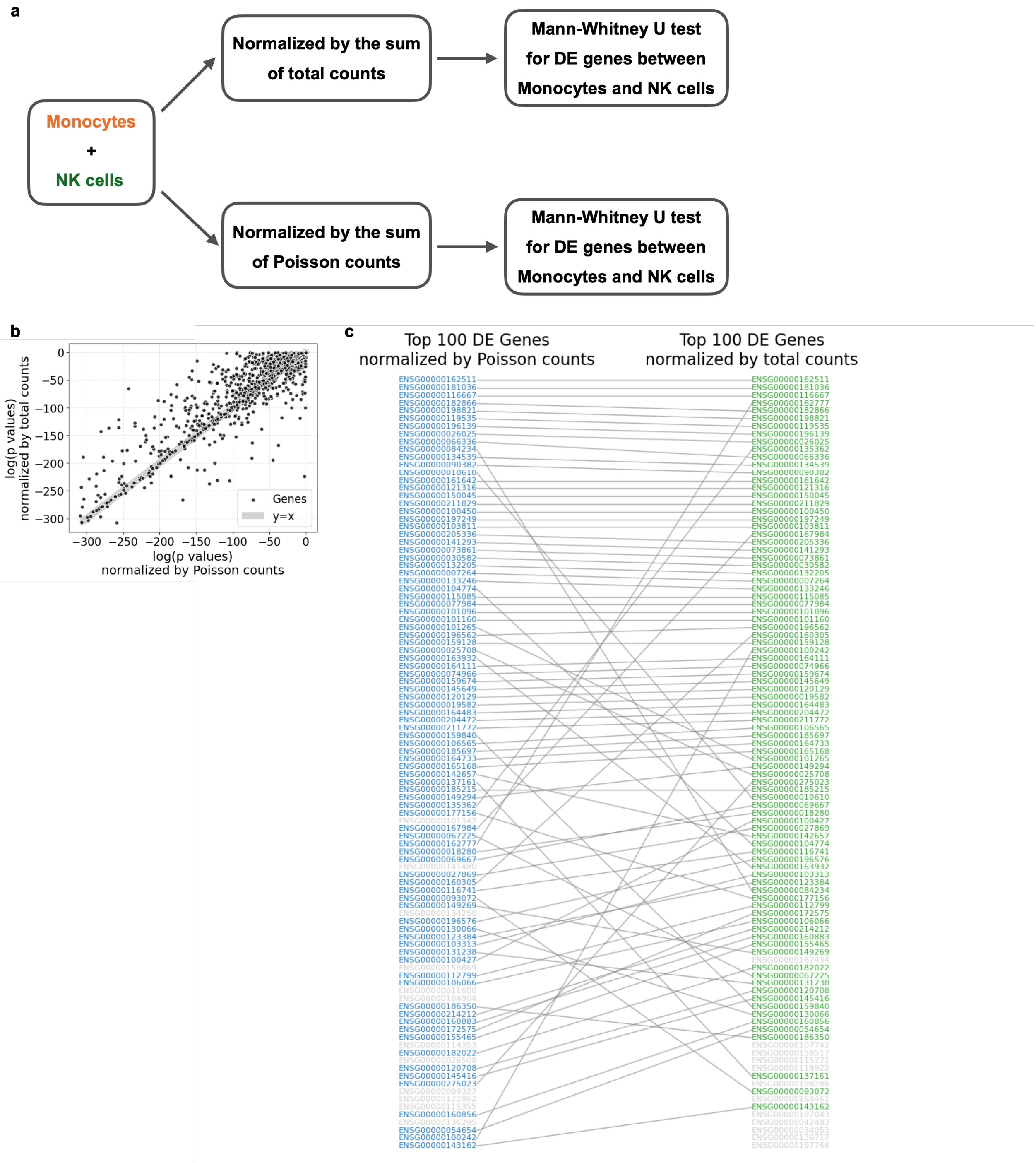


**Figure S8: Supplementary figures for PBMC dataset a)** Procedure for comparing differential expression (DE) analysis results using different cell sizes. **b)** P-values from the Mann–Whitney U test comparing gene expression between monocytes and NK cells, computed after normalizing the data using different cell size estimates. **c)** Correspondence between top 100 genes after two normalizations.

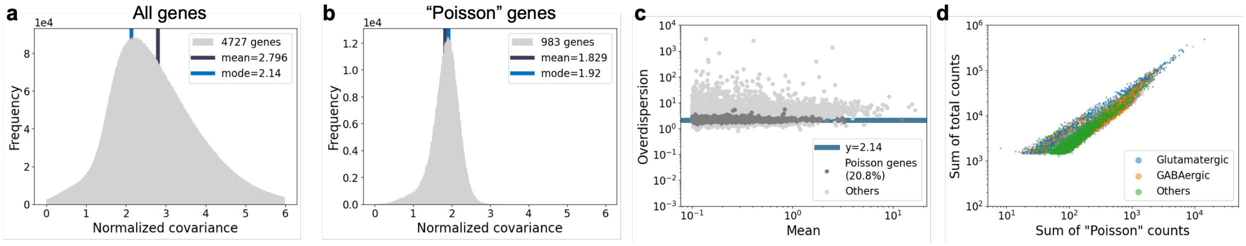


**Figure S9: Results for mouse forebrain dataset(a)** Distribution of normalized covariance between genes with mean expression greater than 0.1 for the PBMC dataset. **(b)** Distribution of normalized covariance between selected Poisson genes for the PBMC dataset. **(c)** Overdispersion-mean relationship for the PBMC dataset. **(d)** Sum of total counts against sum of Poisson counts, colored by cell types.
